# Testing an Optimally Weighted Combination of Common and/or Rare Variants with Multiple Traits

**DOI:** 10.1101/281832

**Authors:** Zhenchuan Wang, Qiuying Sha, Kui Zhang, Shuanglin Zhang

## Abstract

Joint analysis of multiple traits has recently become popular since it can increase statistical power to detect genetic variants and there is increasing evidence showing that pleiotropy is a widespread phenomenon in complex diseases. Currently, most of existing methods test the association between multiple traits and a single common variant. However, the variant-by-variant methods for common variant association studies may not be optimal for rare variant association studies due to the allelic heterogeneity as well as the extreme rarity of individual variants. In this article, we developed a statistical method by testing an optimally weighted combination of variants with multiple traits (TOWmuT) to test the association between multiple traits and a weighted combination of variants (rare and/or common) in a genomic region. TOWmuT is robust to the directions of effects of causal variants and is applicable to different types of traits. Using extensive simulation studies, we compared the performance of TOWmuT with the following five existing methods: gene association with multiple traits (GAMuT), multiple sequence kernel association test (MSKAT), adaptive weighting reverse regression (AWRR), single-TOW, and MANOVA. Our results showed that, in all of the simulation scenarios, TOWmuT has correct type I error rates and is consistently more powerful than the other five tests. We also illustrated the usefulness of TOWmuT by analyzing a whole-genome genotyping data from a lung function study.

## Introductions

Many large cohort studies collected many correlated traits that can reflect underlying mechanism of complex diseases. For example, the UK10K cohort study collected 64 correlated phenotypic traits (The UK10K Consortium *et al.* 2015). Usually complex diseases are characterized by multiple endophenotypes. For example, hypertension can be characterized by systolic and diastolic blood pressure (Newton-Cheh *et al.*2009); metabolic syndrome is evaluated by four component traits: high-density lipoprotein (HDL) cholesterol, plasma glucose and Type 2 diabetes, abdominal obesity, and diastolic blood pressure (Zabaneh and Balding 2010); and schizophrenia can be diagnosed by eight neurocognitive domains (Gur *et al.* 2007). Multiple correlated traits can be influenced by a gene simultaneously. Therefore, by joint analysis of multiple traits, we can not only gain more statistical power to detect pleiotropic variants (Yang and Wang 2012; Solovieff *et al.* 2013; Stephens 2013; Zhou and Stephens 2014; Zhu *et al.* 2015a; Liang *et al.* 2016; Wang *et al.* 2016a; Wang *et al.* 2016b), but also can be important to understand the genetic architecture of the disease of interest (Aschard *et al.* 2014).

Several statistical methods have been developed for testing the association between multiple traits and a single common variant. These methods can be roughly divided into three groups: dimension reduction methods (Klei *et al.* 2008; Ferreira and Purcell 2009; Aschard *et al.* 2014; Wang *et al.* 2016a), regression methods (Korte *et al.* 2012; O’Reilly *et al.* 2012; Zhang *et al.* 2014), and combining test statistics from univariate analysis (O’Brien 1984; Yang *et al.* 2010; van der Sluis *et al.* 2013; Kim *et al.* 2015; Zhu *et al.* 2015b; Liang *et al.* 2016). However, due to the allelic heterogeneity as well as the extreme rarity of rare variants (Li and Leal 2008), the variant-by-variant methods for common variant association studies may not be optimal for rare variant association studies. Recent studies show that complex diseases are caused by both common and rare variants (Pritchard 2001; Pritchard and Cox 2002; Walsh and King 2007; Bodmer and Bonilla 2008; Stratton and Rahman 2008; Kang *et al.* 2010; Teer and Mullikin 2010). Next-generation sequencing technology allows sequencing of the whole genome of large groups of individuals, and thus makes rare variant association studies feasible (Andres *et al.*2007; Metzker 2010). Recently, statistical methods for rare variant association studies with a single trait have been developed by summarizing genotype information from multiple variants. These methods include burden tests (Morgenthaler and Thilly 2007; Li and Leal 2008; Madsen and Browning 2009; Price *et al.* 2010; Zawistowski *et al.* 2010), quadratic tests (Neale *et al.* 2011; Wu *et al.* 2011; Sha *et al.* 2012; Yang *et al.* 2017), and combined tests (Derkach *et al.* 2013; Lee *et al.* 2013; Sha and Zhang 2014; Greco *et al.* 2015).

As we pointed out above, it is essential to develop statistical methods to test the association between multiple traits and multiple variants (common and/or rare variants). Very recently, a few statistical methods for this purpose are appeared (Casale *et al.* 2015; Wang *et al.* 2015; Broadaway *et al.* 2016; Sun *et al.* 2016; Wang *et al.* 2016b; Wu and Pankow 2016). Casale et al. [2015] proposed a set-based association test based on the linear mixed-model. This method enables jointly analyzing multiple correlated traits in rare variant association studies while accounting for population structure and relatedness. Wang et al. [2015] proposed a multivariate functional linear model approach to test association between multiple traits and rare variants in a genomic region. In this approach, the genetic effects of variants are treated as smooth functions of genomic positions of these variants.Gene association with multiple traits (GAMuT) proposed by Broadaway et al. [2016] provide a nonparametric test of independence between a set of traits and a set of genetic variants. This method compares the similarities of multiple traits with the similarities of genotypes at variants in a genomic region. Multivariate Rare-Variant Association Test (MURAT) proposed by Sun et al. [2016] tests association between multiple correlated quantitative traits and a set of rare variants based on a linear mixed model. This method assumes that the effects of the variants follow a multivariate normal distribution with a zero mean and a specific covariance structure. Wu and Pankow [2016] extended the commonly used sequence kernel association test (SKAT) (Wu *et al.* 2011) for a single trait to multiple traits and proposed multiple sequence kernel association test (MSKAT). Wang et al. [2016b] proposed an adaptive weighting reverse regression (AWRR) method. This method uses the score test based on the reverse regression, in which the summation of adaptively weighted genotypes is treated as the response variable and multiple traits are treated as independent variables.

In this article, we developed a new statistical method by testing an optimally weighted combination of variants with multiple traits (TOWmuT) to test the association between multiple traits and a weighted combination of variants (rare and/or common) in a genomic region. TOWmuT is based on the score test under a linear model, in which the weighted combination of variants is treated as the response variable and multiple traits including covariates are treated as independent variables. The statistic of TOWmuT is the maximum of the score test statistic over weights. The weights at which the score test statistic reaches its maximum are called the optimal weights. TOWmuT is applicable to different types of traits and can include covariates. Using extensive simulation studies, we compared the performance of TOWmuT with single-TOW (Sha *et al.* 2012), GAMuT (Broadaway *et al.* 2016), MSKAT (Wu and Pankow 2016), AWRR (Wang *et al.* 2016b) and MANOVA (Yang and Wang 2012). Our results showed that, in all the simulation scenarios, TOWmuT is either the most powerful test or comparable to the most powerful test among the six tests. We also illustrated the usefulness of TOWmuT by analyzing a real whole-genome genotyping data from a lung function study.

## Methods

We consider a sample with *n* unrelated individuals. Each individual has *K* potentially correlated quantitative or qualitative traits (1 for cases and 0 for controls for a qualitative trait) and has been genotyped at *M* variants in a genomic region. Let 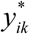 denote the *k* ^*th*^ trait value of the *i*^*th*^ individual and 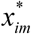 denote the genotype score of the *i*^*th*^ individual at the *m*^*th*^variant, where 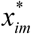 is the number of minor alleles that *i*^*th*^ the individual carries at the *m*^*th*^ variant. We first centralize 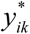 and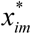 as 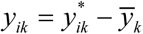 and 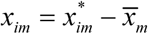 where 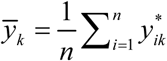 and 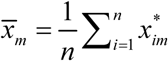. Let *Y*_*i*_ = (*y*_*i1*_,…,*y*_*iK*_)^*T*^, *X*_*i*_ = (*x*_*i1*_,…,*x*_*iM*_)^*T*^, *Y* = (*Y*_*1*_,…,*Y*_*n*_)^*T*^, *X* = (*X*_*1*_,…,*X*_*n*_)^*T*^. For the *i*^*th*^ individual, we consider a linear combination of the variants 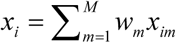 where *w* = (*w*_1_,…,*w*_*M*_)^*T*^ are weights and their values will be decided later.

### Without covariates

We first describe our method without covariates. Consider the linear model

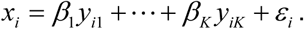

The score test statistic to test the null hypothesis *H*_0_ : *β*_1_ = …= *β*_*K*_ = 0 is given by

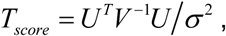

where 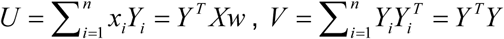 and 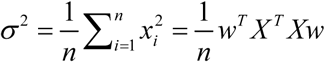 We use 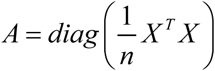 to replace 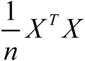 Then *σ* ^2^ becomes 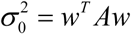 *Aw* and *T*_*score*_ becomes 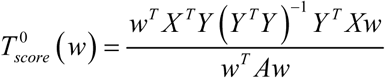. We define the test statistic of TOWmuT as

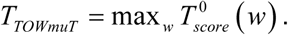

Let *W* = *A*^1/2^*w*, then 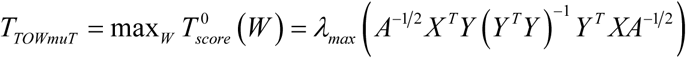, where λ *max* (·) indicates the largest eigenvalue of a matrix. Let *W* ^0^ denote the eigenvector of *A*^-1/2^ *X* ^*T*^*Y* (*Y* ^*T*^*Y*) ^-1^ *Y* ^*T*^ *XA*^-1/2^ corresponding to the largest eigenvalue,*w*^0^ = *A*-1 2*W* 0 is the optimal weights. Actually, we do not need to calculate *w*^0^ in order to calculate *T* _*TOWmuT*_. If we let *C* = *XA*^-1^ *X* ^*T*^, then

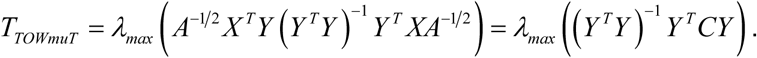

We use a permutation test to evaluate the p-value of *T*_*TOWmuT*_. In details, we randomly shuffle the traits in each permutation. Note that *C* and (*Y* ^*T*^*Y*) ^-1^ do not change in each permutation. Suppose that we perform *B* times of permutations. Let 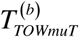 denote the value of *T*_*TOWmuT*_based on the *b*^*th*^ permuted data, where *b* = 0 represents the original data. Then, the p-value of *T*_*TOWmuT*_ is given by

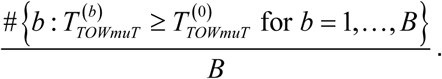

### With covariates

Assume that there are *p* covariates and *z*_*i*1_, …, _*zip*_ individual. Consider the linear model denote the *p* covariates of the *i*^*th*^ In the appendix, we showed that under model

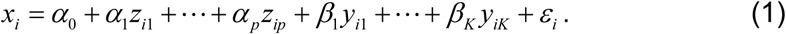

In the appendix, we showed that under model (1), the score test statistic with covariates to test the null hypothesis *H*_0_ : *β*_1_ = … = *β*_*K*_ = 0 is given by

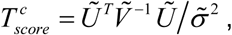

where 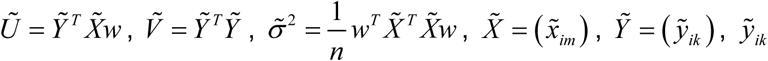 and 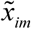 denote the residuals of *y*_*ik*_ and *x*_*im*_ under

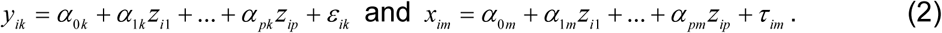

We can see the score test statistic with covariates

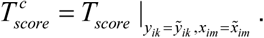

That is, replacing *y*_*ik*_ and *x*_*im*_ by their residuals 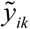 and 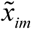 in the score test statistic without covariates *T* _*score*_, it becomes the score test statistic with covariates 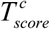.

Therefore, we define TOWmuT statistic with covariates as

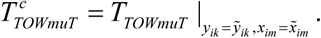

In summary, to apply TOWmuT with covariates, we adjust both trait value *y*_*ik*_ and genotypic score *x*_*im*_ for the covariates by applying linear regressions in (2) and apply TOWmuT without covariates to the residuals 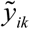 and 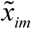

### Comparison of Methods

We compare the performance of our method with the following methods: Multivariate Analysis of Variance (MANOVA) (Liang *et al.* 2016), MSKAT (Wu and Pankow 2016), GAMuT (Broadaway *et al.* 2016), AWRR (Wang *et al.* 2016b) and single-TOW (Sha *et al.* 2012). Here we briefly introduce each of those methods using the notations in the method section.

**MANOVA:** Consider a multivariate multiple linear regression model:

*Y* = *X β* + *ε*, where *Y* denotes the *n* x *K* matrix of phenotypes; *X* denotes the *n* x *M* matrix of genotypes; *β* is a *M* x *K* matrix of coefficients; *ε* is the *n* x *K* matrix of random errors with each row of *ε* to be i.i.d.*MVN* (0, ∑), where ∑ is the covariance matrix of *ε*. To test *H*_0_ : *β* = 0, the likelihood ratio test is equivalent to the Wilk’s Lambda test statistic of MANOVA, that is,

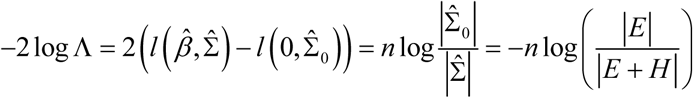.Here Λ denote the ratio of the likelihood function under *H*_0_ to the likelihood function under *H*_1_, *l* (*β*, ∑) is the log-likelihood function,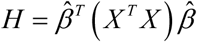 and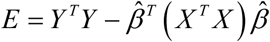 where 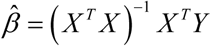 is the maximum likelihood estimator (MLE) of *β*, and ∣ ·∣denotes the determinant of a matrix. The test statistic has an asymptotic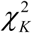 distribution.

**MSKAT**: MSKAT extends the commonly used SKAT (Wu *et al.* 2011) for single trait analysis to test for the joint association of rare variant set with multiple continuous traits.

**GAMuT**: GAMuT compares the similarity in multivariate phenotypes to the similarity in rare-variant genotypes in a genomic region by a machine-learning framework called kernel distance covariance.

**AWRR**: by collapsing genotypes using adaptive weights, AWRR uses the score test to test association based on the reverse regression, in which collapsed genotypes are treated as the response variable and multiple traits are treated as independent variables.

**Single-TOW**: Let 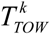 denote the test statistic of TOW to test the association between the *k* th trait and the genotypes at the variants in a genomic region. The test statistic of single-TOW is given by *T*_*single*-*TOW*_ = min_1≤*k* ≤*K*_ *p*_*k*_, where *p*_*k*_ is the p-value of 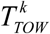 for *k* = 1, …, *K*. The p-value of *T*_*single*-*TOW*_ is estimated using a permutation procedure.

### Simulations

In our simulation studies, we use the empirical Mini-Exome genotype data provided by the genetic analysis workshop 17 (GAW17) to generate genotypes. This dataset contains genotypes of 697 unrelated individuals on 3205 genes. We choose four genes: ELAVL4 (gene1), MSH4 (gene2), PDE4B (gene3), and ADAMTS4 (gene4) with 10, 20, 30, and 40 variants, respectively. Then, we merge the four genes to form a super gene (Sgene) with 100 variants. In our simulation studies, we generate genotypes based on the genotypes of 697 individuals in the Sgene because the distribution of the minor allele frequencies (MAFs) in the Sgene can represent the distribution of MAFs in all of the 3205 genes (Sha *et al.* 2012). To generate a qualitative disease affection status, we use a liability threshold model based on a continuous phenotype (quantitative trait). An individual is defined as affected if the individual’s phenotype is at least one standard deviation larger than the phenotypic mean. This yields a prevalence of 16% for the simulated disease in the general population. In the following, we describe how to generate a quantitative trait.

We consider that all causal variants are rare (MAF < 0.01). We randomly choose *n*_*c*_ rare variants as causal variants, where *n*_*c*_ is determined by the percentage of causal variants among rare variants. We use *n*_*r*_ and *n*_*p*_ to denote the number of risk rare variants and protective rare variants, respectively, where *n* _*r*_ + *n* _*p*_ = *n* _*c*_. Let 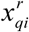 and 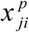 denote the genotypic scores of the *q*^*th*^ risk rare variant and the *j*^*th*^ protective rare variant for the *i*^*th*^ individual, respectively. We assume that genotypes impact on *L* traits. Let *h* and *h*_*l*_ denote the heritability of all the *nc* rare causal variants for the *L* traits and the *l*^*th*^ trait among the *L* traits, respectively. We generate *L* random numbers *t*_1_, …, _*tL*_ from a uniform distribution between 0 and 1. *L* Then, the heritability of *l*^*th*^ trait among the *L* traits is 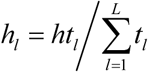. Given the heritability of the *l*^*th*^ trait *h*_*l*_, we generate *n*_*c*_ random numbers 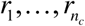 from a uniform distribution between 0 and 1. The heritability of the *m*^*th*^ causal variant for the *l*^*t*^ trait is given by 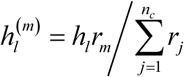

In our simulation studies, we consider two covariates *Z*_1_ and *Z* _2_, where *Z*_1_ is a continuous covariate generated from a standard normal distribution, and *Z* _2_ is a binary covariate taking values 0 and 1 with a probability of 0.5. We generate *K* traits by considering the factor model (van der Sluis *et al.* 2013; Aschard *et al.* 2014; Wang *et al.* 2016a)

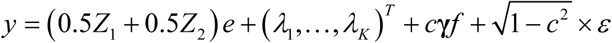

where *y* = (*y*_1_, …, *y*_*k*_) ^*T*^; = (1, …, 1) ^*T*^;λ =(*λ* _1_, …, *λ*_*k*_) is the vector involved genotypes;*f* = (*f*_1_, …, *f*_2_) ^*T*^∼ *MVN* (0, ∑), ∑ = (1 - *ρ*) *I* + *ρA*, *A* is a matrix with elements of 1, *I* is the identity matrix, and *ρ* is the correlation between *f*_*i*_ and *f* _*j*_; *R* is the number of factors; **γ** is a *K* by *R* matrix; *c* is a constant number; *ε* = (*ε* _1_, …, *ε* _*T*_) ^*T*^ a vector of residuals; and *ε* _1_, …, *ε* _*K*_ are independent, *ε* _*k*_ ∼ *N* (0,1) for *k* = 1, …, *K*.

We consider the following six models with different number of factors and different number of traits affected by genotypes. In these models, the within-factor 1correlation is *c*^2^ and the between-factor correlation is *ρ* = *ρc*^2^.

**Model 1:** There is only one factor and genotypes impact on 6 traits with the same effect size. This is equivalent to set *R* = 1 and **γ** = (1, …, 1) ^*T*^. In details,

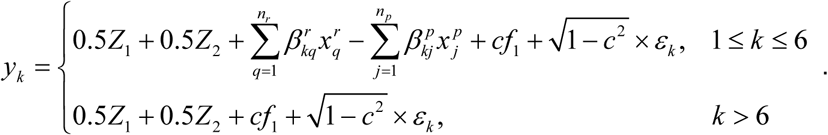

**Model 2:** There are five factors and genotypes impact on 6 traits. We set *R* = 5 and **γ** = *diag* (*D*_1_, *D*_2_,*D*_*3*_, *D*_*4*_, *D*_*5*_), where 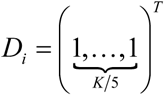 for *i* = 1, …,5. In details,

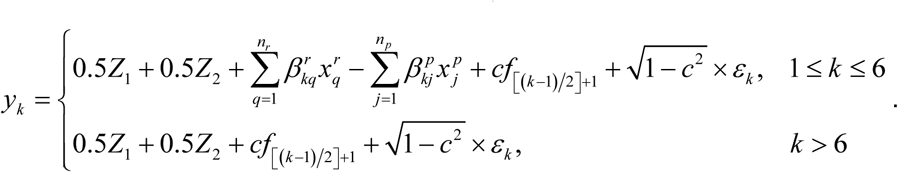

**Model 3:** There are two factors and genotypes impact on 6 traits. That is,*R* = 2 and **γ** = *diag* (*D*_1_, *D*_2_), where 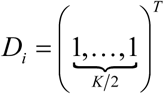 for *i* = 1,2. In details,

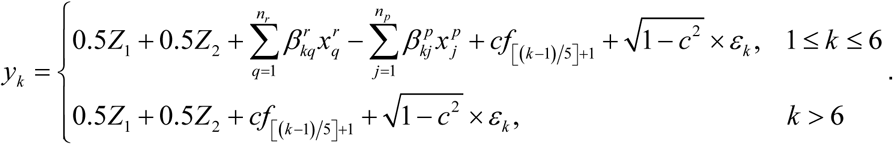

**Model 4:** There are five factors and genotypes impact on one trait. That is,*R* = 5 and **γ** = *diag* (*D*_1_, *D*_2_,*D*_*3*_, *D*_*4*_, *D*_*5*_), where 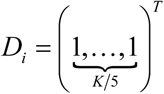 for *i* = 1,…,5. In details,

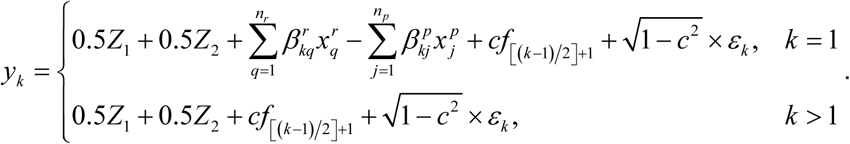

**Model 5:** There are only two factors and genotypes impact on one trait. That is, *R* = 2 and **γ** = *diag* (*D*_1_, *D*_2_), where 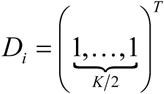 for *i* = 1,2. In details,

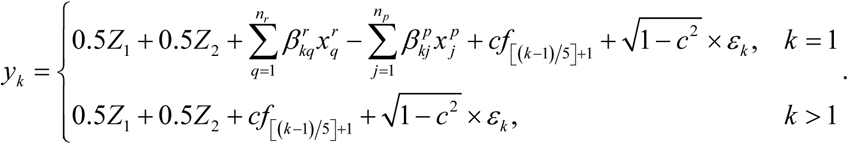

**Model 6:** There is *K* factors and genotypes impact on 6 traits. That is, *R* = *K*,**γ=***I* and *c* = 1. In details,

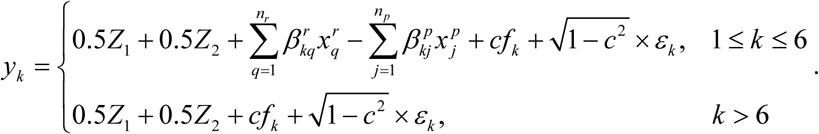

## Results

To evaluate the type I error rates of the proposed test TOWmuT, we set *λ*_*k*_ = 0 for *k* = 1, …, *K* in the 6 models. We consider different sample sizes, different significance levels, different models, and different types of traits. In our simulations we consider 10 traits (K = 10). In each simulation scenario, the p-values of TOWmuT are estimated by 1000 permutations and the type I error rates of TOWmuT are evaluated using 10,000 replicated samples. For 10,000 replicated samples, the 95% confidence intervals (CIs) for the estimated type I error rates of nominal levels 0.05 and 0.01 are (0.046, 0.054) and (0.008, 0.012), respectively. The estimated type I error rates of TOWmuT are summarized in Tables 1 and 2. From these two tables, we can see that 70 out of 72 (greater than 95%) estimated type I error rates are within the 95% CIs and the two estimated type I error rates not within the 95% Cis (0.05555 and 0.01295) are very close to the bound of the corresponding 95% CI, which indicates that TOWmuT is valid.

**Table 1.** The estimated type I error rates of TOWmuT for 10 quantitative traits under each model with covariates.

|  | Sample Size |  |  |  |
| --- | --- | --- | --- | --- |
|  | Model | 500 | 1000 | 2000 |
| $\alpha = 0.05$ | 1 | 0.05365 | 0.0515 | 0.0515 |
|  | 2 | 0.0521 | 0.0528 | 0.0504 |
|  | 3 | 0.0513 | 0.0540 | 0.0503 |
|  | 4 | 0.0514 | 0.0511 | 0.05 |
|  | 5 | 0.05381 | 0.04825 | 0.05 |
|  | 6 | 0.0482 | 0.0508 | 0.05325 |
| $\alpha = 0.01$ | 1 | 0.01165 | 0.0098 | 0.0117 |
|  | 2 | 0.012 | 0.01015 | 0.0102 |
|  | 3 | 0.01175 | 0.01075 | 0.0113 |
|  | 4 | 0.01145 | 0.01075 | 0.0118 |
|  | 5 | 0.01141 | 0.01095 | 0.0117 |
|  | 6 | 0.0097 | 0.0105 | 0.01185 |

**Table 2.** The estimated type I error rates of TOWmuT for the mixture of five quantitative traits and five qualitative traits under each model with covariates.

|  | Sample Size |  |  |  |
| --- | --- | --- | --- | --- |
|  | Model | 500 | 1000 | 2000 |
| $\alpha = 0.05$ | 1 | 0.05365 | 0.05385 | 0.05005 |
|  | 2 | 0.0511 | 0.0483 | 0.05115 |
|  | 3 | 0.0508 | 0.05375 | 0.052 |
|  | 4 | 0.0529 | 0.04915 | 0.0536 |
|  | 5 | 0.054 | 0.05355 | 0.04825 |
|  | 6 | 0.05555 | 0.0493 | 0.0529 |
| $\alpha = 0.01$ | 1 | 0.0105 | 0.01295 | 0.00995 |
|  | 2 | 0.0105 | 0.009 | 0.0097 |
|  | 3 | 0.01145 | 0.0104 | 0.0101 |
|  | 4 | 0.01065 | 0.00945 | 0.01165 |
|  | 5 | 0.0118 | 0.0105 | 0.00875 |
|  | 6 | 0.01195 | 0.00935 | 0.01105 |

For power comparisons, we consider different values of heritability, different models, different types of traits, different percentages of protective variants, different values of between-factor correlation, and different values of within-factor correlation. In each of the simulation scenarios, the p-values of TOWmuT, AWRR and single-TOW are estimated using 1,000 permutations and the p-values of MANOVA, GAMuT, and MSKAT are estimated using asymptotic distributions. The powers of all of the six tests are evaluated using 1,000 replicated samples at a significance level of 0.05.

Figure 1 gives the power comparisons of the six tests (Single-TOW, MSKAT, AWRR, MANOVA, GAMuT, and TOWmuT) for the power as a function of the total heritability based on the six models for 10 quantitative traits. This figure shows that (1) TOWmuT is consistently the most powerful one among the six tests; (2) MANOVA is the second most powerful when genotypes impact on multiple traits (models 1-3 and 6) while AWRR is the second most powerful when genotypes impact on a single trait (models 4-5); (3) MSKAT is consistently less powerful than other multivariate tests probably because SKAT gives larger weights than that of TOW to only those variants with MAF in the range (0.01,0.035) and there are only 8% variants with MAF in the range (0.01,0.035) in Sgene which our simulations are based on; and (4) MSKAT and GAMuT have similar powers in all six models.

**Figure 1.**
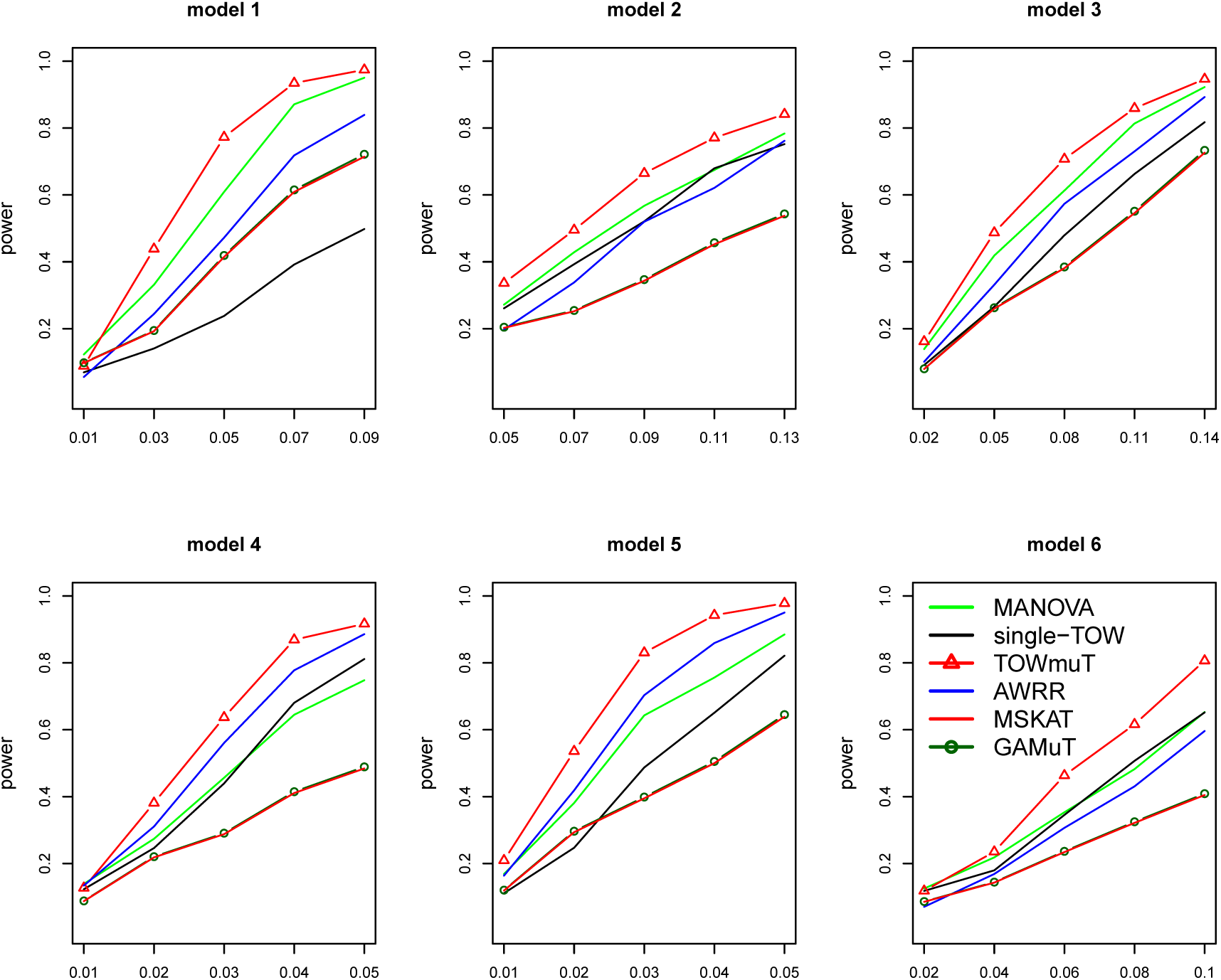
Power comparisons of the six tests (Single-TOW, MSKAT, AWRR, MANOVA, GAMuT and TOWmuT) for the power as a function of total heritability for 10 quantitative traits with covariates. The sample size is 1000. The between-factor correlation is 0.3 and the within-factor correlation is 0.7. The percentage of the causal variants is 0.2. All causal variants are risk variants.

Figure 2 gives the power comparisons of the five tests (Single-TOW, AWRR, MSKAT, GAMuT, and TOWmuT) for the power as a function of the total heritability for the mixture of 5 quantitative traits and 5 qualitative traits. We only compare the powers of five tests because MANOVA has inflated type I error rate in this case. This figure shows that (1) TOWmuT is consistently the most powerful one among the five tests; (2) AWRR is second most powerful when genotypes impact on multiple traits (models 1-3 and 6) while MSKAT and GAMuT are second most powerful when genotypes impact on a single trait (models 4-5); (3) MSKAT and GAMuT have similar powers in all six models; and (4) single-TOW is consistently less powerful than other four multivariate tests because we keep correlations between traits similar to that in Figure 1 such that correlations between original quantitative traits are larger than that in Figure 1.

**Figure 2.**
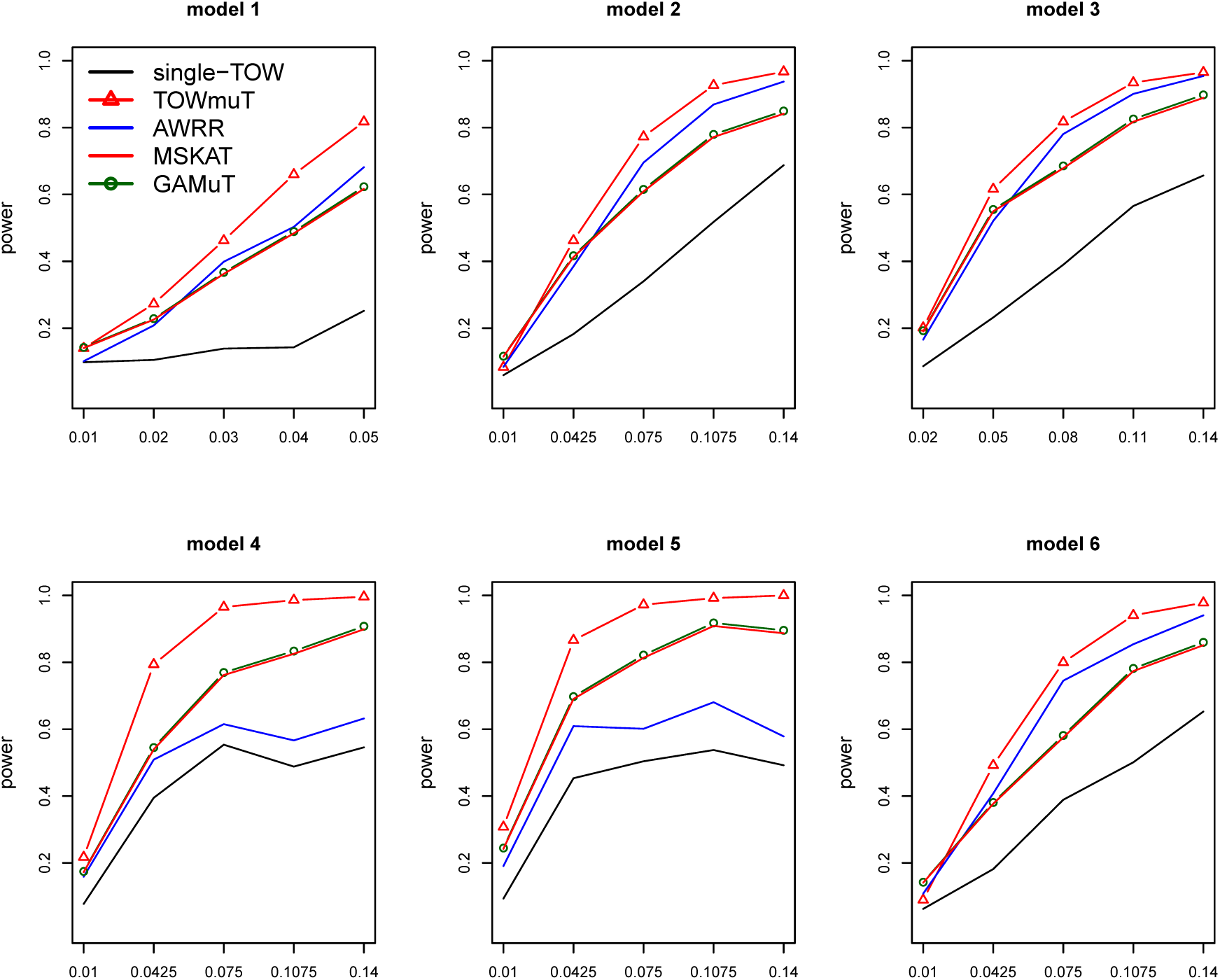
Power comparisons of the five tests (Single-TOW, AWRR, GAMuT, MSKAT and TOWmuT) for the power as a function of heritability for the mixture of half quantitative traits and half qualitative traits with covariates. The sample size is 1000. Covariance matrix of 10 traits is similar to that of 10 quantitative traits with between-factor correlation being 0.3 and the within-factor correlation being 0.7. The percentage of the causal variants is 0.2. All causal variants are risk variants.

We also compare the powers of the six tests for the power as a function of the within-factor correlation for models 1-5 and between-factor correlation for model 6 for 10 quantitative traits (Figure S1). As shown in this figure, the power of single-TOW is robust to the between-factor correlation or the within-factor correlation since the minimum p-value-based approach is largely unaffected by the trait correlation (Wu and Pankow 2016). However, with the increasing of the between-factor correlation or within-factor correlation, the power of other five tests essentially increases. Other patterns of the power comparisons are similar to those of in Figure 1.

Power comparisons of the six tests for the power as a function of the percentage of protective variants for 10 quantitative traits are given by Figure S2. This figure shows that the power of all six tests are robust to the percentage of protective variants, therefore, all of these methods are robust to the directions of the genetic effects. Other patterns of the power comparisons are similar to those of in Figure 1.

### Application to the COPDGene

Chronic obstructive pulmonary disease (COPD) is a common disease in elderly patients that causes significant morbidity and mortality (Nazir and Erbland 2009). The Genetic Epidemiology of COPD Study (COPDGene) (Regan *et al.* 2010) was designed to identify genetic factors associated with COPD. In this COPDGene study, a total of more than 10,000 subjects have been enrolled including 2/3 non-Hispanic Whites (NHW) and 1/3 African-Americans (AA). In this analysis, we only include 5,430 NHW with no missing phenotypes. Each of the 5,430 NHW has been genotyped at 630,860 SNPs. Based on the literature studies of COPD (Han *et al.*2011; Chu *et al.* 2014; Liang *et al.* 2016), we selected 7 key quantitative COPD-related phenotypes, including FEV1 (% predicted FEV1), Emphysema (Emph), Emphysema Distribution (EmphDist), Gas Trapping (GasTrap), Airway Wall Area (Pi10), Exacerbation frequency (ExacerFreq), Six-minute walk distance (6MWD), and 4 covariates, including BMI, Age, Pack-Years (PackYear) and Sex.

To evaluate the performance of our proposed method on a real data set, we applied six methods (TOWmuT, MANOVA, MSKAT, GAMuT, AWRR, and single-TOW) to the COPDGene of NHW population to test the association between each of 50-SNP blocks and the 7 key quantitative COPD-related phenotypes. To identify significant 50-SNP blocks associated with the phenotypes, we used Bonferroni correction to decide the significance level. The total number of 50-SNP blocks is 12617, therefore, the Bonferroni corrected significance level is 0.05/12617 ≈4×10^−6^. Table 3 summarized the significant blocks identified by at least one method. There were total six significant blocks in Table 3. All of the six blocks have been previously reported to be in association with COPD or lung functions (Pillai *et al.* 2009; Cho *et al.* 2010; Figarska *et al.* 2014; Lutz *et al.* 2015). PDSS1 and ABI1 are located between LOC107984176 and LOC105376467, which are Intergenic regions and contain the SNPs associated with pulmonary function (Imboden *et al.* 2012; Lutz *et al.* 2015). From Table 3, we can see that TOWmuT identified four blocks; AWRR identified two blocks; MANOVA, MSKAT and GAMuT identified one block; single-TOW did not identify any blocks. From these results, we can see that TOWmuT identified the most of significant 50-SNP blocks among the six methods, which is consistent with the results of our simulation studies.

**Table 3.**
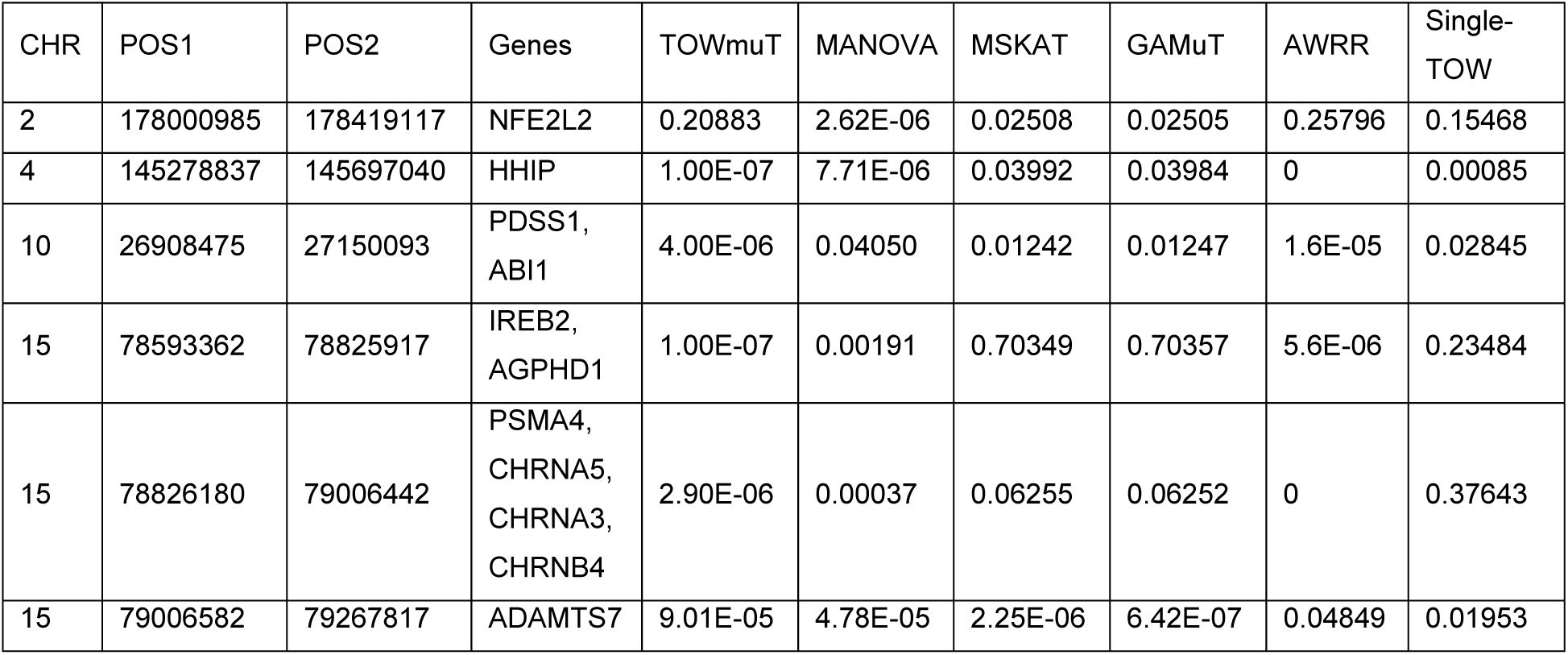
Significant blocks identified by at least one method (p-values less than 4×10^−6^) and the corresponding p-values in the analysis of COPDGene.

## Discussion

We developed TOWmuT to perform joint analysis of multiple traits in gene-based association studies based on the following reasons: (1) multiple related traits are usually measured in genetic association studies of complex diseases; (2) there is increasing evidence showing that pleiotropy is a widespread phenomenon in complex diseases; and (3) there is a shortage of gene-based approaches for multiple traits. We used extensive simulation studies to compare the performance of TOWmuT with MANOVA, MSKAT, AWRR, GAMuT and Single-TOW. Our results showed that TOWmuT has correct type I error rates and is consistently more powerful than other five methods. Additionally, the real data analysis results demonstrated that the proposed method has great potential in gene-based association study for complex diseases with multiple phenotypes such as COPD.

Recently, it has become a major focus of investigation to identify a small number of rare causal variants that contribute to complex diseases (Capanu and Ionita-Laza 2015). Several methods to pinpoint the causal variants have been developed for testing the association with a single trait. These methods include backward elimination (BE) method (Ionita-Laza *et al.* 2014), hierarchical model method (Ionita-Laza *et al.* 2014), and adaptive combination of p-values method (Lin 2016). To extend the TOWmuT method to identify a small number of causal variants which are associated with multiple traits, we can use the BE method. In each step, we remove one variant that has the smallest contribution to the association between multiple traits and the set of variants and then we evaluate the p-value for testing association between multiple traits and the remaining variants by TOWmuT. Causal variants are the set of variants corresponding to the smallest p-value.

The computation time required for running TOWmuT depends on the sample size, the number of variants in the genomic region, the number of traits, and the number of permutations. The running time of TOWmuT with 1000 permutations on a data set with 5000 individuals, 7 traits and 10 variants in a genomic region on a laptop with 4 Intel Cores @ 3.30GHz and 4 GB memory is about 0.14s. To perform genome-wide association studies, we can first select genomic regions that show evidence of association based on a small number of permutations (e.g. 1,000), and then a large number of permutations are used to test the selected regions.

## Acknowledgements

Research reported in this publication was supported by the National Human Genome Research Institute of the National Institutes of Health under Award Number R15HG008209. The content is solely the responsibility of the authors and does not necessarily represent the official views of the National Institutes of Health.

The Genetic Analysis workshops are supported by NIH grant R01 GM031575 from the National Institute of General Medical Sciences. Preparation of the Genetic Analysis Workshop 17 Simulated Exome Data Set was supported in part by NIH R01 MH059490 and used sequencing data from the 1000 Genomes Project (www.1000genomes.org).

This research used data generated by the COPDGene study, which was supported by NIH grants U01HL089856 and U01HL089897. The COPDGene project is also supported by the COPD Foundation through contributions made by an Industry Advisory Board comprised of Pfizer, AstraZeneca, Boehringer Ingelheim, Novartis, and Sunovion.

Superior, a high-performance computing infrastructure at Michigan Technological University, was used in obtaining results presented in this publication.

The authors have no conflict of interests to declare.

## Appendix

We use the same notations in the method section. Let *Y* = (*Y*_1_,… *Y*_*n*_) ^*T*^, *Z*_*i*_=(1,*Z*_i1_,…, *Z*_*ip*_) ^*T*^ *Z* = (*Z*_*1*_, …, *Z*_*n*_) ^*T*^, and *x* = (*x*_1_, *…, x*_*n*_) ^*T*^Under the linear model

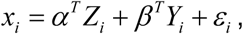

the log-likelihood (up to a constant) is given by

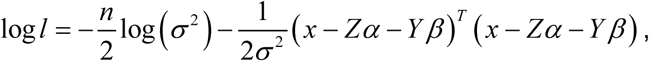

where *α* = (*α*_0_, …, *α* _*p*_) ^*T*^, *β* = (*β*_1_, …, *β*_*k*_) ^*T*^, and *ε* _1_, …, *ε n* are independent and *ε* _*I*_ ∼ *N* (0,*σ* ^2^) Then,

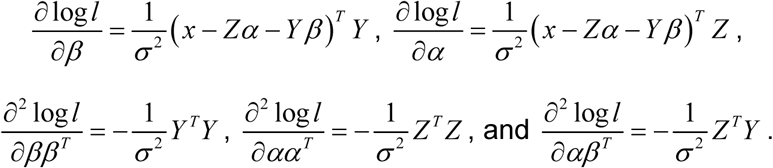

Let 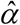 and 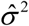 denote the maximum likelihood estimates of *α* and*σ* ^2^ under null hypothesis *H*_0_: *β* = 0. Then, 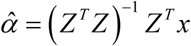 and 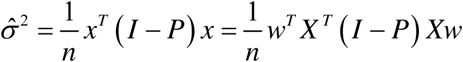, where *P* = *Z* (*Z*^*T*^ *Z*) ^T^ *Z*^*T*^. Let *θ* = (*α* ^*T*^, *β* ^*T*^) ^*T*^. The score and information matrix are 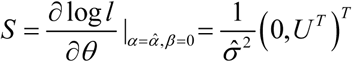 and 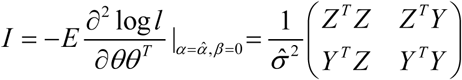, where *U* = *Y* ^*T*^(*I* - *P*) *x*= *Y* ^*T*^(*I* - *P*) *Xw*. The score test statistic is given by

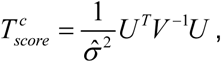

where *V* = *Y* ^*T*^(*I* - *P*) *Y*. Note that (*I* - *P*)^2^ = *I* - *P*. We have 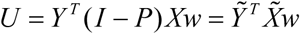, 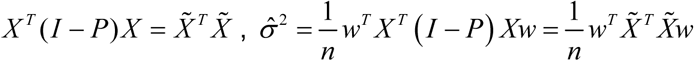, and 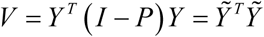 where 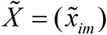 and 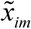 is the residual of *x*_*im*_ under the linear regression model (2); 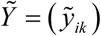 and 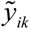 is the residual of *y*_*ik*_ under the linear regression model (2).

Therefore,

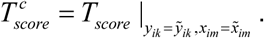

